# Differential Privacy Protection Against Membership Inference Attack on Machine Learning for Genomic Data

**DOI:** 10.1101/2020.08.03.235416

**Authors:** Junjie Chen, Wendy Hui Wang, Xinghua Shi

**Author notes:** © 2016 The Authors. Open Access chapter published by World Scientific Publishing Company and distributed under the terms of the Creative Commons Attribution Non-Commercial (CC BY-NC) 4.0 License.

## Abstract

Machine learning is powerful to model massive genomic data while genome privacy is a growing concern. Studies have shown that not only the raw data but also the trained model can potentially infringe genome privacy. An example is the membership inference attack (MIA), by which the adversary, who only queries a given target model without knowing its internal parameters, can determine whether a specific record was included in the training dataset of the target model. Differential privacy (DP) has been used to defend against MIA with rigorous privacy guarantee. In this paper, we investigate the vulnerability of machine learning against MIA on genomic data, and evaluate the effectiveness of using DP as a defense mechanism. We consider two widely-used machine learning models, namely Lasso and convolutional neural network (CNN), as the target model. We study the trade-off between the defense power against MIA and the prediction accuracy of the target model under various privacy settings of DP. Our results show that the relationship between the privacy budget and target model accuracy can be modeled as a log-like curve, thus a smaller privacy budget provides stronger privacy guarantee with the cost of losing more model accuracy. We also investigate the effect of model sparsity on model vulnerability against MIA. Our results demonstrate that in addition to prevent overfitting, model sparsity can work together with DP to significantly mitigate the risk of MIA.

## 1. Introduction

Genomics has emerged a frontier of data analytics empowered by machine learning and deep learning, thanks to the rapid growth of genomic data that contains individual-level sequences or genotypes at large scale. It is critical to collect, aggregate, and share to build powerful and robust machine learning models for genomics analysis. However, genetic privacy is a growing and legitimate concern that prevents wide sharing of genomic data. Genomic data is naturally sensitive and private, and the sharing of such data can potentially disclose an individual’s sensitive information such as identity, disease susceptibility or family history.^1,2^ Regulations and rules, such as Health Insurance Portability and Accountability Act (HIPAA)^3^ have been established to address the usage and disclosure of individuals’ health information for covered entities. However, HIPAA does not cover the protection of de-identified genomic data, or genomic data generated from entities which are not covered by HIPAA such as commercial sequencing or genotyping service providers.

In addition to imposing regulations, rules, ethnics (e.g. consent forms), common practices of privacy protection of genomic data, such as the controlled access of individual-level genomic data (e.g. the database of Genotypes and Phenotypes or dbGaP^4^), are thus limited. Intensive investigation should be carried out to develop robust technologies to protect genetic privacy while allowing for wide sharing of genomic data. Toward an overaching goal of achieving trustworthy biomedical data sharing and analysis, we are in great need of new techniques for genetic privacy protection including the design and development of computational strategies to mitigate the following two types of the leakage of genetic privacy:

- *Privacy leakage via sharing data*: the individual records may be leaked by sharing raw genomic data or summary statistics data; and
- *Privacy leakage via sharing models*: sharing machine learning models built upon genomic analysis may leak information about an individual’s genomic data in the training dataset.^5^

While most of the prior works focus on the former type of privacy leakage resulted from sharing data,^6–9^ in this study, we mainly focus on the latter type of privacy leakage resulted from sharing machine learning models. Although less explored, this later type of privacy leakage does pose an emerging risk in genomics study given the increasing application and sharing of machine learning models in various genomic analysis scenarios. One particular scenario that is of broad interest in agriculture, animal breeding, and biomedical science, is a conventional problem of genomic selection or phenotype prediction. Phenotype prediction refers to the investigation of using genetic features (e.g. genetic variants) to predict phenotype labels, which relies heavily on statistical and machine learning models applied to high-dimensional genomic data. Hence, in this study, we focus on the evaluation of privacy leakage and protection based on machine learning models for phenotype prediction, using two demonstrative strategies. These two demonstrative cases include a classical sparse learning model (e.g. Lasso) and a promising deep learning model such as convolutional neural networks (CNNs).

Although there exists a wide spectrum of attacks on machine learning models, the *membership inference attack* (MIA)^10^ has recently attracted research efforts for introducing privacy leakage when sharing machine learning models. More specifically, MIA refers to an attack to infer if the target record was included in the model’s training dataset where machine learning models are shared (not the training data though). MIA has been demonstrated as an effective attack on images and relational data (e.g.,^5,10,11^). However, it remains unclear if MIA is effective on genomic data which significantly differ from conventional imaging or relational data. Therefore, we will investigate the efficiency of MIA on machine learning models for phenotype prediction based on genomic data.

To defend against various attacks (including MIA), a few techniques have been developed to mitigate privacy leakage, including homomorphic encryption,^12^ federated learning,^13^ and differential privacy (DP).^8^ While homomorphic encryption and federated learning are mainly used to provide privacy protection for data sharing DP provides a popular solution for publicly sharing information not only about the data but also the models.^14^ Recently, multiple defense mechanisms against MIA^15–17^ have been explored, with DP^18^ standing out as an efficient strategy that provides a rigorous privacy guarantee against MIA.^10^ Previous studies on imaging data^19,20^ have shown that DP is an effective solution for granting wider access to machine learning models and results while keep them private. Therefore, we will mainly consider DP as a defense mechanism against MIA, given its theoretical privacy guarantee and its applicability to protecting protection for data and models.

Hence, in this study, we investigate the effectiveness of using DP as a defense mechanism to prevent the risk of sharing two widely used machine learning technologies (e.g. Lasso and CNN), against MIA for phenotype prediction on genomic data. The **main contributions** of our paper are in two folds:

### First

we investigate the vulnerability of machine learning against MIA on genomic data, and evaluate the effectiveness of using DP as a defense mechanism. Although MIA and DP have been respectively studied in other areas including imaging and databases, they are rarely explored in genomic studies especially when they are combined together as a dual strategy for attack and defense as in this study. Particularly, we evaluate the trade-off between the defense power against MIA and the prediction accuracy of the target model under various privacy settings of DP. Our results show that the relationship between the privacy budget and target model accuracy can be modeled as a log-like curve, and hence there exists a trade-off between privacy and accuracy near the inflection point. These results point to invaluable insights and potential directions in future studies to comprehensively evaluate how to choose the privacy budget of DP against MIA that can best address the trade-off between privacy and model accuracy, which is a challenging balance to achieve in real-case scenarios.

### Second

we investigate the effect of model sparsity on privacy vulnerability to and defense effectiveness against MIA. Genomic data is primarily high dimensional, where the feature size is significantly larger than sample size. Hence, adding sparsity (e.g. the regularization terms in Lasso models) to machine learning models on genomic data is a critical and effective strategy to alleviate the curse of dimensionality and avoid overfitting. Our results show that model sparsity together with DP can significantly mitigate the risk of MIA, in addition to providing robust and effective models. Other than genomic data, these findings may be applied to other types of high dimensional data, and model sparsity should be investigated together with DP as a powerful mechanism for defending against attacks on machine learning models.

## 2. Related Work

### Membership inference attack (MIA)

Shokri *et al.*^10^ is the first work that defines MIA and inspires a few follow-up studies. For example, Truex *et al.*^21^ characterize the attack vulnerability with respect to the types of learning models, data distribution, and transferablity. Salem *et al.*^5^ design new variants of MIA by relaxing the assumptions of model types and data. Long *et al.*^11^ generalize MIA by identifying vulnerable records and indirect inference. Liu *et al.*,^22^ Song *et al.*^23^ and Hayes *et al.*^24^ propose new MIA variants against deep learning models including variational autoencoders (VAEs) and generative adversarial networks (GANs).

### Differential privacy (DP)

DP^14^ has become the most widely used approach that measures the disclosure of privacy pertaining to individuals. Various DP mechanisms have been developed^25,26^ including a DP algorithm for logistic regression^27^ and a random forest algorithm with DP.^28^ Going beyond classic machine learning models, Shokri *et al.*^29^ adapted DP to deep neural networks. Abadi *et al.*^20^ developed a differentially private stochastic gradient descent (SGD) algorithm for the TensorFlow framework.

### Privacy-preserving genomic data analysis

The leakage of genetic privacy has been frequently reported where the identity or sensitive trait of a particular individual was disclosed by the identity tracing attacks,^30,31^ attribute disclosure attacks via DNA,^4^ and completion attacks.^32,33^ Several privacy-preserving solutions for genomic data analytics have been proposed, including DP strategies for genome-wide association studies (GWAS),^7,34^ crytographic solutions including homomorphic encryption^12,35^ and secure multi-party computation (MPC) solutions.^35^ However, these privacy-preserving methods aim to mitigate privacy leakage when data is shared and none of them have considered privacy leakage when sharing models.

## 3. Methods

In this section, we introduce the methods used in our study, including differential privacy (DP), and membership inference attack (MIA). The supplementary materials and source code are available at https://github.com/shilab/DP-MIA.git.

### Membership Inference Attack (MIA)

MIA is a privacy-leakage attack that predicts whether a given record was used in training a target model based on the output of the target model for the given record.^29^ As illustrated in **Fig. 1**, MIA assumes that a target machine learning model is trained on a set of labeled data records sampled from a certain population. Then the adversary utilizes the output of the target model of a given record or sample to infer the membership of the given record (i.e., the given record was included in the training dataset of the target model). Formally, let *f_target_*() be the target model trained on a private dataset 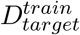 which contains labeled data records (**x**, *y*). The output of the target model is a probability vector **y** = *f_target_*(**x**) whose size is the number of classes that label the given records. Let *f_shadow_* () be the attack model trained on a dataset 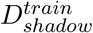, which is generated by the attacker to mimic the training data of the target model. We use the same assumption as in the pioneering work^10^ that the shadow dataset is disjoint from the private target dataset used to train the target model (i.e., 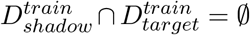). The shadow model *f_shadow_*() aims to mimic the performance of the target model *f_target_*() (i.e. take similar inputs and outputs of the target model). Let *f_attack_*() be the attack model. Its input **x**_attack_ is composed of a predicted probability vector and a true label, where the distribution of predicted probability vectors heavily depends on the true label. Since the goal of the attack is membership inference, the attack model is a binary classifier, in which the output 1 indicates that the target record is in the training dataset, and 0 otherwise.

**Fig. 1.**
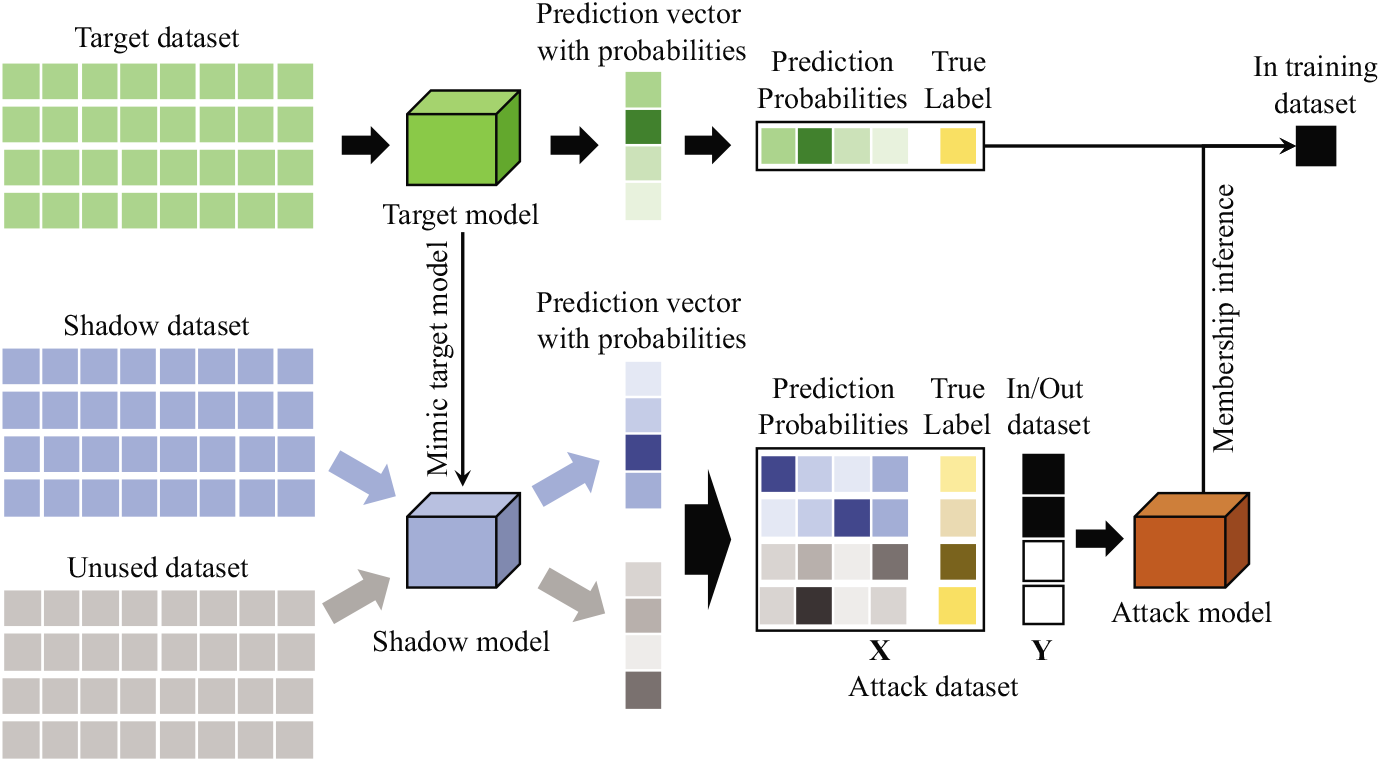
An illustration of membership inference attack.

To construct the MIA model, a shadow training technique is often applied to generate the ground truth of membership inference. One or multiple shadow models are built to imitate the target model. While previous works^10^ mainly consider the black-box access to the target model, we consider the white-box setting, where the adversary has the full knowledge of the target model, including its hyperparameters, network structure, and activation functions used at each layer. This white-box threat model reflects the observations that in collaborative learning, researchers share their full models as well as the fact that white-box representations of models may fall into the hands of an adversary via other means (e.g., a security breach). With the white-box access, the shadow model *f_shadow_*() has the same architecture as the target model *f_target_*(). Consequently, the key problem of MIA is to train the shadow models on the shadow dataset that has the same format and distribution as the target dataset.

### Differential privacy (DP)

DP describes the statistics of groups while withholding individuals’ information within the dataset.^14^ Informally, DP ensures that the outcome of any data analysis on two databases differing in a single record does not vary much. Formally, a randomized algorithm 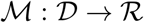 with domain 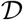 and range 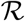 is (*ε, δ*)-differentially private if for all subsets of 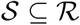 and for all database inputs 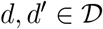 such that ||*d* ‒ *d*′||_1_ ≤ 1 satisfied with 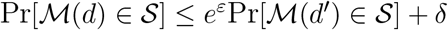, where ||*d* – *d*′||_1_ requires that the number of records that differ between *d* and *d*′ is at most 1. The parameter *ε* is called the *privacy budget.* Intuitively, lower *ε* indicates stronger privacy protection. If *δ* = 0, we say 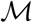 is plain *ε*-differentially private, and simplify (*ε*, 0)-differential privacy as *ε*-differential privacy. The parameter *δ* controls the probability that plain *ε*-differential privacy is violated. A lower *δ* value signifies greater confidence of differential privacy. A rule of thumb of values of *δ* is to be smaller than the inverse of the training data size 1/||*d*||.^20^

## 4. Experimental Setup

### 4.1. Dataset

We evaluate effectiveness of DP against MIA on a widely-used yeast genomic dataset.^36^ We choose this yeast dataset because it provides an ideal scenario for evaluating the power and privacy of phenotype prediction with well-controlled genetic background and phenoytpe quantifications. Without worries about complex genetic background and the hard-to-defined phenotypes in humans, our usage of this simple case of yeast data will allow us to directly evaluate the performance of MIA, DP and machine learning models. We extracted and filtered the missing values of the original data^36^ and organized the dataset in a matrix that contains genotypes of 28,820 genetic variants or features (with values of 1 and 2 representing the allele comes from crossing from a laboratory strain or a vineyard strain respectively) form 4,390 individuals. The dataset also contains the phenotypes of these 4,390 individuals for 20 end-point growth traits,^36^ where we pick Copper Sulfate for our target phenotype or label in this use case. Hence, similar to any typical human genomic data, the yeast data is high dimensional where the feature size (28,820) is much larger than sample size (4,390). Since MIA is mainly launched on classification models, in this study, we binarize the quantitative phenotype values (i.e. Copper Sulfate measurements) as 1 if they are larger than the mean value, and 0 otherwise.

### 4.2. Implementation of Target Models

We implement a Lasso model and a CNN model as the target model, as an illustration of a widely used machine learning and deep learning model on high-dimensional genomics data.

#### Least absolute shrinkage and selection operator (Lasso)

Lasso is a regression analysis method that performs variable selection with *ℓ*_1_ norm regularization in order to avoid overfitting.^37^ Due its good interpretability, Lasso has been a widely used baseline for association analysis in many genetic analysis task (e.g. GWAS,^38^ epistasis,^39^ eQTL^40^). Lasso minimizes the residual sum of squares subject to the sum of the absolute value of the coefficients being less than a constant. The general objective of Lasso is 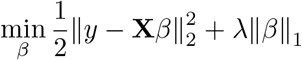, where **X** is the genotype matrix, *β* is coefficient vector, and *y* is the gene expression trait vector. λ is the coefficient of *ℓ*_1_ norm which controls the model sparsity. Lasso uses an Ą norm regularization to shrink the parameters of the majority of features to zero which are trivial, and those variants corresponding to non-zero terms are selected as the identified important features. We set values of λ as 0 (without model sparsity) and 0.001352 (with model sparsity selected using glmnet package in R^41^).

#### Convolutional neural network (CNN)

CNN has shown its capability to capture the spatial or local patterns in genomic data, such as data correlated patterns including linkage disequilibrium in genotypes.^42,43^ For demonstration, the CNN model in this study is designed with one CNN layer, followed by a dense layer as an output layer. To improve model robustness, the *ℓ*_1_ norm is applied to all layers to shrink small weights to zeros. We utilize a grid search with 5-fold cross validation to find the optimized hyperparameters. In particular, we use two different learning rates, 0.01 and 0.001. We use two micro batches of size 50% and 100% of batch size. Regarding *ℓ*_2_ norm clipping which determines the maximum amounts of *ℓ*_2_ norm clipped to cumulative gradient across all network parameters from each microbatch, we use four unique *ℓ*_2_ norm clipping values, 0.6, 1.0, 1.4, and 1.8. We use two different kernel sizes, 5 and 9, for CNN models. The number of kernels is in {8,16}. Furthermore, we set the coefficient λ of *ℓ*_2_ norm to achieve the sparsity of CNN models. We set the value of λ as 0 (without model sparsity) and 0.001352 (with model sparsity chosen using glmnet^41^).

### 4.3. Implementation of Differential Privacy

We implement DP on both Lasso and CNN models with and without *ℓ*_1_ norm respectively, using a Python library called TensorFlow-privacy,^44^ which implements DP by adding standard Gaussian noise on each gradient on the SGD optimizer. The major process for training a model with parameters *θ* by minimizing the empirical loss function *L*(*θ*) with differentially private SGD, is shown as the following: at each step of computing the SGD: 1) compute the gradient ∇_θ_*L*(*θ,x_i_*) for a random subset of examples; 2) clip the *ℓ*_2_ norm of each gradient; 3) compute the average of gradients; 4) add some noise in order to protect privacy; 5) take a step in the opposite direction of this average noisy gradient; 6) in addition to outputting the model, compute the privacy loss of the mechanism based on the information maintained by the privacy accountant.

In the implementation, the privacy budget is determined by a function that takes multiple hyperparameters as the input. These hyperparameters include the number of epochs, batch size and the noise multiplier. The noise multiplier controls the amount of noises added during training. In general, adding more noise leads to better privacy and lower utility. The privacy budgets and their corresponding hyperparameters used in this study are as follows (details in **Table S.1** of the supplementary materials): the number of epochs ∈ {50,100}, the batch size ∈ {8,16} and the noise multiplier ∈ {0.4,0.6,0.8,1.0,1.2}. We set the value of the parameter *δ* as the inverse of training dataset size (i.e. *δ* = 0.00066489).^20^

### 4.4. Implementation of Membership Inference Attack

We split the whole dataset into two disjoint subsets, one as the private target dataset and the other one as the public shadow dataset for training differentially private machine learning models and performing MIA correspondingly.^10^ We randomly split the public shadow dataset, with 80% used for model training and 20% used to generate the ground truth of the attack model. We focus on a white-box model attack, where the target model’s architecture and weights are accessible. Thus the shadow model has the same architecture and hyperparameters as the target model to check how much privacy will be leaked in the worst case. We use an open-source library of MIA^45^ to conduct MIA attacks on the Lasso and CNN models. We build one shadow model on the shadow dataset to mimic the target model, and generate the ground truth to train the attack model. The attack dataset is constructed by concatenating the probability vector output from the shadow model and true labels. If a sample is used to train the shadow model, the corresponding concatenated input for the attack dataset is labeled ‘in’, and ‘out’ otherwise. For the attack model, we build a random forest with 10 estimators and a max depth of 2. Each MIA attack is randomly repeated 5 times.

### 4.5. Evaluation Metrics

Our evaluation metrics include: (1) the mean accuracy of 5-fold cross validation of the target model on the private target dataset, and (2) the mean of MIA accuracy of 5 MIA attacks. The accuracy of the target model on the training (testing, resp.) data is measured as the precision (i.e., the fraction of classification results that are correct) of the prediction results on the training (testing, resp.) data. We follow the pioneering work^10^ and use the *attack accuracy* to measure MIA performance. The attack accuracy is calculated as the fraction of records inferred as members are indeed true members of the target dataset.

## 5. Results

### 5.1. Vulnerability of target model against MIA without DP protection

We first investigate the vulnerability of Lasso and CNN models against MIA for predicting the target phenotype without any DP protection. **Table 1** shows the accuracy of the two different target models without DP and attack accuracy of MIA on these models. When the models are not sparse (λ = 0), Lasso and CNN achieves similar accuracy on the target dataset (0.7910 vs. 0.7894). The attack accuracy of MIA on Lasso and CNN with no sparsity is 0.5728 and 0.5726 respectively, which is better than random guess (0.5). After introducing sparsity by adding an *ℓ*_1_ norm (λ = 0.001352) to model coefficients (in Lasso) or weights (in CNN), the target accuracy of either model is slightly improved and their attack accuracy is slightly reduced. These results demonstrate that MIA can be used to infer the membership of particular records when training Lasso and CNN models on genomic data.

**Table 1.** Model performance against MIA (without DP).

| Methods | Target model |  | Attack model |  |
| --- | --- | --- | --- | --- |
|  | Accuracy | Std. | Accuracy | Std. |
| Lasso ($\lambda = 0$) | 0.7910 | 0.0123 | 0.5728 | 0.0071 |
| Lasso ($\lambda = 0.001352$) | 0.7963 | 0.0157 | 0.5631 | 0.0042 |
| CNN ($\lambda = 0$) | 0.7894 | 0.0199 | 0.5726 | 0.0059 |
| CNN ($\lambda = 0.001352$) | 0.7936 | 0.0225 | 0.5628 | 0.0050 |

### 5.2. Impact of privacy budget on target model accuracy

In order to evaluate the impact of DP on the accuracy of the target model, we conduct a grid search to find different privacy budgets and quantitatively investigate the impact of privacy budget on model accuracy as summarized in **Fig. 2(a)**. As expected, we observe that the performance of all target models rapidly deteriorates as the privacy budget becomes smaller. When the privacy budget is large, both non-sparse Lasso (λ = 0) and non-sparse CNN (λ = 0) models achieve similar accuracy performance. Additionally, for sparse Lasso (λ = 0.001352) and sparse CNN (λ = 0.001352) models, their accuracy is much more downgraded by DP compared with the models without DP, even when the privacy budget is large. This is because sparse models only keep the coefficients/weights which are higher than λ, and shrink those coefficients/weights that are smaller than λ to 0. Therefore, adding any noise to those large weights will have more significant impact on the accuracy of the target model.

**Fig. 2.**
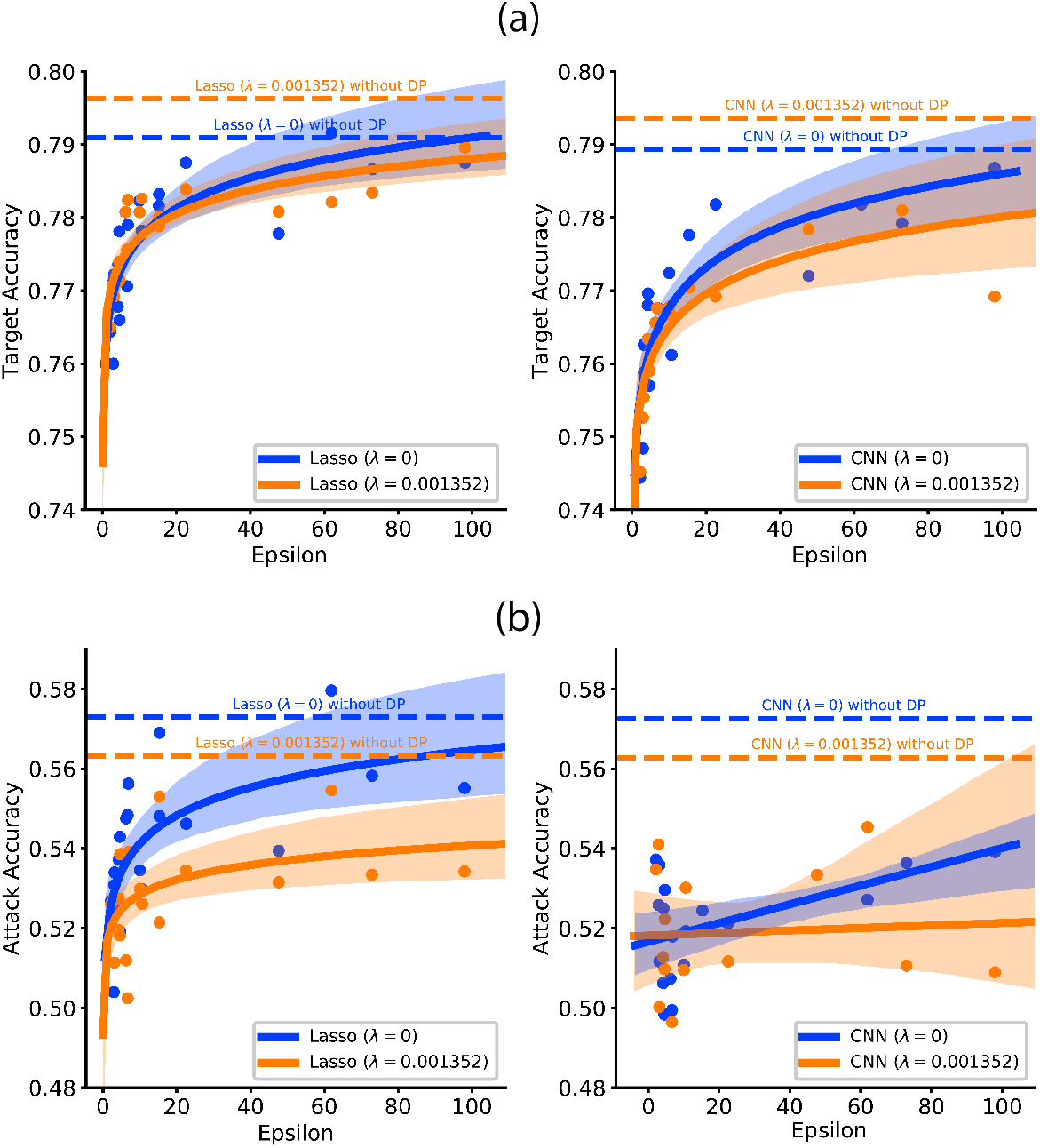
Accuracy values of the (a) target model and (b) attack model respectively under various privacy budgets (5-fold cross validation). Curves indicate the fitted regression lines; shadow areas represent the 95% confidence intervals for corresponding regressions. Horizontal dotted lines represent model performances without DP.

We also observe that there exists a trade-off between privacy and accuracy of target models, which is critical for deciding the balance between privacy and utility of the models. As shown in **Fig. 2(a)**, the fitting curve between the privacy budget and the model accuracy can be represented as a log-like curve. Thus, we can choose the turning point near the inflection points of the curve as the balance between privacy budget and model accuracy. Based on this observation, we can choose the privacy budget of 10 that best addresses the trade-off between privacy and target accuracy in this study.

### 5.3. Effectiveness of DP against MIA

To evaluate the effectiveness of DP against MIA, we conduct MIA on the target models with different DP budgets. Our results (**Fig. 2(b)**) show that DP can defend against MIA effectively. The reason that DP can defend against MIA is because DP perturbs the prediction vector output by the target model, so that the adversary cannot easily infer the membership from such noisy prediction. Furthermore, the attack accuracy of MIA decreases as the privacy budget becomes smaller (i.e., stronger privacy protection). In particular, for Lasso, a smaller privacy budget (*ε* ≤ 10) rapidly reduces the attack accuracy. But when the privacy budget *ε* exceeds 10, the attack accuracy stays relatively stable. For CNN, the attack accuracy keeps steadily decreasing when ε increases.

### 5.4. Effect of model sparsity

We investigate the effect of model sparsity by adding an *ℓ*_1_ norm to the weights. Due to the large hyperparameter searching space, we only use the value of λ = 0.001352 for both Lasso and CNN, which is chosen by using the glmnet package in R.^41^ λ = 0 means the model has no sparsity. Our results (**Table 1**) show that model sparsity can improve the accuracy of the target model and reduce the attack accuracy of MIA when DP is not deployed. This is because on the high-dimensional dataset, a Lasso or CNN model with no sparsity (i.e. λ = 0) can overfit the training data. However, by introducing model sparsity, the overfitting of the model is reduced, leading to better accuracy of the target model.

We further explore the impact of sparsity on the accuracy of the target model when DP is deployed. We observe that sparse models have slightly worse target model accuracy under different privacy budgets (**Fig. 2(a)**). This is because each weight in a sparse model is important to prediction results; and any perturbation to these weights can significantly impact model accuracy. We also find that when the privacy budget is smaller than the trade-off (e.g. *ε* < 10 in our results), the accuracy of the target model is relatively insensitive to model sparsity compared with larger privacy budgets (i.e., *ε* > 10). Next, we evaluate the impact of model sparsity on the defense power of DP against MIA. As shown in **Fig. 2(b)**, sparse models provide better privacy protection compared with those models without sparsity, given the same DP budget *ε*. Our results demonstrate that a privacy budget of 10 best addresses the trade-off between privacy and defense against MIA on the dataset under investigation.

## 6. Conclusion

We investigate the vulnerability of trained machine learning models for phenotype prediction on genomic data against a new type of privacy attack named membership inference attack (MIA), and evaluate the effectiveness of using differential privacy (DP) as a defense mechanism against MIA. We find the MIA can successfully infer if a particular individual is included in the training dataset for both Lasso and CNN models, and DP can defend against MIA on genomic data effectively with a cost of reducing accuracy of the target model. We also evaluate the trade-off between privacy protection against MIA and the prediction accuracy of the target model. Moreover, we observe that introducing sparsity into the target model can further defend against MIA in addition to implementing the DP strategy.

Using yeast genomic data as a demonstration, our study provides a novel computational framework that allows for investigating not only the privacy leakage induced from MIA attacks on machine learning models, but also the efficiency of classical defending mechanisms like DP against these new attacks.. In the future, we will apply this framework to large-scale human genomic data to further investigate the impact of DP against MIA in human genomic analysis. As the extension of our white-box attack model, we will consider the black-box access of MIA on the target model, where the adversary simply uses the target model as a black-box for query but does not have any inside information of the model. We will also investigate other factors (e.g., the number of classes) and other types of genomic analysis (e.g. associations studies, risk prediction) to comprehensively assess the attack power of MIA and the effectiveness of appropriate defense mechanisms. In this study, we evaluate the efficiency of DP against MIA on Lasso and CNN based models, we plan to investigate other machine learning models and deep architectures including GAN and its variants.

## 7. Acknowledgement

Preprint of an article submitted for consideration in Pacific Symposium on Biocomputing (©[2021] World Scientific Publishing Co., Singapore, http://psb.stanford.edu/.

