## Supplemental material for "Differential Privacy Protection Against Membership Inference Attack on Machine Learning for Genomic Data"

### S.1. Figures

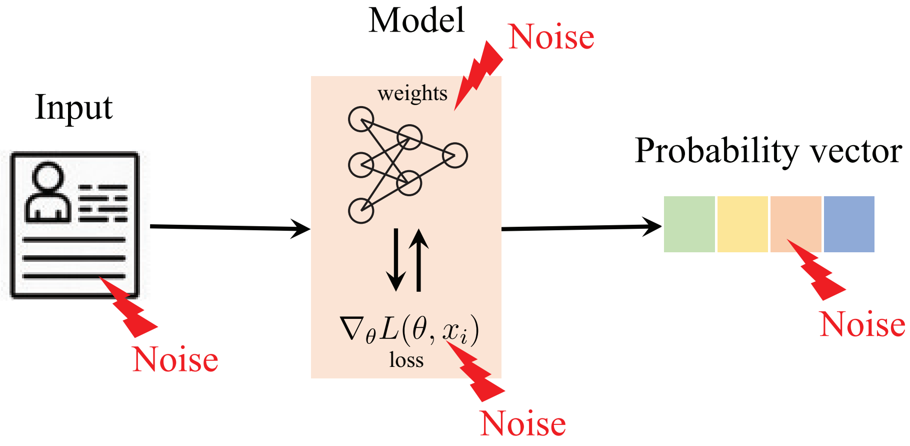


Fig. S.1. An illustration of four types of methods to implement differentiallly private machine learning models. DP can be implemented by adding noises to the input data, to the final weights, to loss gradient and to output probabilities.

### S.2. Tables

Table S.1. The epsilon $\varepsilon$ value as a function of epoch $\in \{50, 100\}$, batch $\in \{8, 16\}$ and noise multiplier $\in \{0.4, 0.6, 0.8, 1.0, 1.2\}$, under the condition of $\delta = 0.00066489$, using differentially private SGD. Larger epoch and batch result in larger epsilon, while larger noise multiplier result in small epsilon.

| epsilon | epoch | batch | noise multiplier |
|---|---|---|---|
| 2.1602 | 50 | 8 | 1.2 |
| 2.9187 | 50 | 8 | 1.0 |
| 3.1102 | 100 | 8 | 1.2 |
| 3.1596 | 50 | 16 | 1.2 |
| 4.2119 | 100 | 8 | 1.0 |
| 4.3068 | 50 | 16 | 1.0 |
| 4.5779 | 100 | 16 | 1.2 |
| 4.6433 | 50 | 8 | 0.8 |
| 6.2594 | 100 | 16 | 1.0 |
| 6.6847 | 100 | 8 | 0.8 |
| 6.8959 | 50 | 16 | 0.8 |
| 10.0391 | 100 | 16 | 0.8 |
| 10.6203 | 50 | 8 | 0.6 |
| 15.2897 | 100 | 8 | 0.6 |
| 15.3320 | 50 | 16 | 0.6 |
| 22.5690 | 100 | 16 | 0.6 |
| 47.6580 | 50 | 8 | 0.4 |
| 61.9543 | 50 | 16 | 0.4 |
| 72.9451 | 100 | 8 | 0.4 |
| 98.0023 | 100 | 16 | 0.4 |
